# Natural Colonization of Laboratory Mice with *Staphylococcus aureus* Primes a Systemic Immune Response

**DOI:** 10.1101/114314

**Authors:** Daniel Schulz, Dorothee Grumann, Patricia Trübe, Kathleen Pritchett-Corning, Sarah Johnson, Kevin Reppschläger, Janine Gumz, Nandakumar Sundaramoorthy, Stephan Michalik, Sabine Berg, Jens van den Brandt, Richard Fister, Stefan Monecke, Benedict Uy, Frank Schmidt, Barbara M. Bröker, Siouxsie Wiles, Silva Holtfreter

## Abstract

**Background:** Whether mice are an appropriate model for *S*. *aureus* infection and vaccination studies is a matter of debate, because they are not considered as natural hosts of *S. aureus.* Sparked by an outbreak of S. *aureus* infections in laboratory mice, we investigated whether laboratory mice are commonly colonized with S. *aureus* and whether this might impact on infection experiments.

**Methods:** We characterized 99 *S. aureus* isolates from laboratory mice (*spa* typing, virulence gene PCR), and quantified murine antibodies using FlexMap technology.

**Results:** Specific-pathogen-free mice from various vendors were frequently colonized with *S. aureus* (0-21%). *S. aureus* was readily transmitted from murine parents to offspring, which became persistently colonized. Most murine isolates belonged to the lineage CC88 (54%). Murine strains showed features of host adaptation, such as absence of *hlb*-converting phages and superantigen genes, as well as enhanced coagulation of murine plasma. Importantly, *S. aureus* colonization induced a systemic IgG response specific for numerous *S. aureus* proteins, including several vaccine candidates.

**Conclusion:** Laboratory mice are natural hosts of *S. aureus* and, therefore, provide better infection models than previously assumed. Pre-exposure to S. *aureus* is a possible confounder in *S. aureus* infection and vaccination studies.

## Background

*Staphylococcus aureus* is a dangerous opportunistic bacterial pathogen, a leading cause of hospital and community infections worldwide, and a prominent example of the antibiotic resistance crisis [1]. There is currently no vaccine available [2]. Thus, novel approaches for the prevention and treatment of infections are urgently required. Mice are the most common surrogate host to model *S. aureus* infection with the advantages of having a well-characterized immune system, many gene knock-out strains available, and being relatively easy and inexpensive to breed. However, whether mice are appropriate has often been questioned because there is broad consensus in the research community that mice are not natural hosts of *S. aureus* [3-6]. Moreover, experimental colonization of mice with *S. aureus* is usually transient, and high infection doses are routinely required [7].

Reports on natural *S. aureus* colonization or infection of laboratory mice are scarce [8, 9]. We have reported an outbreak of *S. aureus* infections in mice bred in a university-associated animal facility [10-12]. Male mice suffered from preputial gland adenitis (PGA), which is the most common location for abscesses in mice [8]. The causative strain, JSNZ, belongs to CC88, a lineage rarely found among human and animal isolates [13-16].

Adaptation to new hosts is a complex process, involving the loss and/or acquisition of mobile genetic elements (MGEs), such as phages, plasmids, and pathogenicity islands, as well as the accumulation of mutations in virulence genes resulting in host-specific allelic variants or loss of function [17, 18]. The most prominent example for host adaptation is *hlb*-integrating Sa3int phages. These encode the human-specific immune evasion gene cluster (IEC), which carries genes for staphylokinase (*sak*), staphylococcal complement inhibitor (*scn*) and chemotaxis inhibitory protein of *S. aureus* (CHIPS; *chp*) as well as staphylococcal enterotoxins A or P (sea, *sep*) [19, 20]. These phages are common in human isolates, but frequently lacking in animal-adapted strains, including the mouse-adapted JSNZ [10, 20].

Laboratory animal vendors generally produce mice to two different microbiological quality levels. Specific-pathogen-free (SPF) mice are free of infectious agents that are known to cause illness in mice, interfere with research, or are zoonotic [21]. Specific and opportunistic pathogen free (SOPF) mice are maintained free of additional microbial agents. *S. aureus* is considered an opportunistic pathogen in mice and not routinely excluded from SPF barrier rooms. Since most laboratories use SPF mice, natural *S. aureus* colonization of experimental mice might be more widespread than expected. Since previous exposure of mice to *S. aureus* may influence the results of experimental infection or vaccination, it is important to learn more about *S. aureus* colonization in laboratory mice.

In this study, we analyzed the prevalence of *S. aureus* in SPF mice from all the main vendors, determined whether murine *S. aureus* isolates are adapted to their host, and investigated if naturally colonized mice are primed against *S. aureus.*

## Methods

### Health reports of laboratory mice

Reports were obtained from the official websites of Charles River (http://www.criver.com/products-services/basic-research/health-reports/), The Jackson Laboratories (https://www.jax.org/jax-mice-and-services/customer-support/customer-service/animal-health/health-status-reports), Taconic (http://www.taconic.com/quality/health-reports/), Janvier Labs (http://www.janvier-labs.com/rodent-research-models-services/research-models.html), Envigo (http://www.envigo.com/products-services/research-models-services/resources/health-monitoring-reports/) in November 2016.

### Murine *S. aureus* strains

Charles River's Research Animal Diagnostic Services (Wilmington, USA) provided 99 *S. aureus* isolates (CR strains) from laboratory mice they were sent by customers. In addition, five *S. aureus* strains and sera from colonized healthy C57BL/6 mice and five strains from C57BL/6 mice with preputial gland adenitis (PGA) were obtained from the Charles River breeding facilities at Kingston NY and Hollister CA, respectively. SPF mice and caretakers from an animal facility at the University of Greifswald, Germany, were screened for *S. aureus* colonization.

### Human *S. aureus* strains

Human *S. aureus* strains (n=107) were obtained from healthy blood donors [22, 23]. Human CC88 isolates were obtained from various clinical sources [10, 13] (Table S4).

### Ethics statement

Human plasma samples were obtained from healthy volunteers. All participants were adults and gave written informed consent. The study was approved by the ethics board of the Medical Faculty of the University of Greifswald (BB 014/14; 24.01.2014). Murine blood samples and nasopharyngeal swabs were obtained from C57BL/6 SPF and SOPF mice during routine health monitoring at Charles River facilities in Hollister CA, Kingston NY, and Wilmington MA, USA. The study was approved by the Charles River Institutional Animal Care and Use Committee (Protocol number P06172002 – Holding & Euthanasia of Animals for Diagnostic Testing and Health Monitoring). All animal work was performed following United States Public Health Service Policy on Humane Care and Use of Laboratory Animals and the US Animal Welfare Act.

### *S. aureus* identification and genotyping

*S. aureus* was identified by colony morphology on mannitol salt agar (MSA) plates, *S. aureus-*specific latex agglutination test (Staph Xtra Latex kit, ProLexTM, Richmond Hill, ON, Canada) as well as gyrase and nuclease PCR (see below). *Spa* genotyping and multilocus sequence typing (MLST) were performed as described elsewhere [24, 25].

### Virulence gene detection

Multiplex PCRs were applied to detect 25 *S. aureus* virulence genes, including gyrase (*gyr*), methicillin resistance (*mecA*), Panton-Valentine leukocidin (*pvl*), staphylococcal superantigens (*sea-selu, tst*), exfoliative toxins (*eta, etd*) and *agr* group 1-4 [22, 23]. *S. aureus* bacteriophage types (*Sa1int – Sa7int*) were detected by multiplex PCR as previously reported [22]. The Sa3int-phage-encoded IEC genes were detected by multiplex PCR using primers specific for *sak, chp, scn, Sa3int,* and *gyrase* (see Supplemental Material).

### Coagulation assay

65 μl bacterial stationary cultures were mixed with 500 μl of human or murine heparinized plasma (Equitech-Bio, Kerrville, USA) and incubated at 37 °C without agitation. Coagulation was visually examined in a blinded fashion using a modified coagulation score [26]: no coagulation = 0; small coagulation flakes = 1; medium-sized clot = 2; large clot = 3; complete coagulation (tube can be inverted) = 4.

### Detection of anti-*S. aureus* serum antibodies

Murine serum IgG against extracellular *S. aureus* proteins was determined by indirect ELISA using protein A-deficient *S. aureus* (JSNZ∆*spa*, PGA12∆*spa*). Serum IgG specific to 58 recombinant *S. aureus* proteins was quantified using FLEXMAP 3D^™^ technology [27]; normalized mean fluorescent intensities were calculated as measures of antibody binding intensity.

### Statistics

Data analysis was performed using GraphPad Prism. Antibody titers in colonized, infected and SOPF mice were not normally distributed and therefore compared with the Mann-Whitney-Test, using the Bonferroni correction for multiple testing.

## Results

### *S. aureus* frequently colonizes SPF mice

To investigate the prevalence of *S. aureus* in commercially available laboratory mice, we screened publicly available health reports from the major vendors (Charles River, The Jackson Laboratories (JAX), Taconic, Janvier labs, and Envigo). The highest cumulative rates were reported in SPF mice from the North American Charles River facilities (20.9%), followed by Taconic (9.7% and 8.5% in the US and Europe, respectively) (Table 1). In contrast, standard colonies from JAX were almost free of *S. aureus* (0.8%). Importantly, the European branches of Janvier, Charles River, and Envigo do not report the *S. aureus* status of SPF mice as this is not requested by the Federation of Laboratory Animal Science Associations (FELASA) [21]. At all vendors, SOPF barrier rooms, isolators, and areas housing immune-deficient mice are actively managed to exclude opportunistic pathogens and were *S. aureus-*free.

**Table 1:** Percentage of *S. aureus*-positive mice and barriers in different breeding facilities, sorted by vendor and barrier status (cumulative data extracted from publicly available health reports in November 2016).

| Vendor | Barrier status<sup>1</sup> | % *S. aureus* positive mice (absolute No.)<sup>2</sup> |  | % *S. aureus* positive barriers (absolute No.)<sup>2</sup> |  |
| --- | --- | --- | --- | --- | --- |
| Charles River, Europe | SPF | ND<sup>3</sup> |  | ND<sup>3</sup> |  |
|  | SOPF | 0.0% | (0/6766) | 0.0% | (0/22) |
|  | Immunodeficient | 0.0% | (0/51) | 0.0% | (0/2) |
| Charles River, US and CAN | VAF | 20.9% | (360/1724) | 65.2% | (15/23) |
|  | SOPF (ELITE) | 0.0% | (0/2524) | 0.0% | (0/7) |
|  | Immunodeficient | 0.0% | (0/11565) | 0.0% | (0/5) |
| The Jackson laboratory, US | Standard | 0.8% | (15/1828) | 9.0% | (1/11) |
|  | High | 0.0% | (0/1094) | 0.0% | (0/6) |
|  | Maximum | 1.0% | (160/16140)<sup>4</sup> | 13.8% | (4/29) |
|  | Low barrier (RAF) | 0% | (0/894) | 0.0% | (0/21) |
|  | Intermediate barrier (RAF) | 0% | (0/321) | 0.0% | (0/7) |
|  | Elevated barrier (RAF) | 0% | (0/60) | 0.0% | (0/1) |
| Taconic, Europe | Murine Pathogen Free™ | 8.5% | (42/494) | 25.0% | (2/8) |
|  | Excluded Flora and Restricted Flora™ | 0.0% | (0/1648) | 0.0% | (0/7) |
| Taconic, US | Murine Pathogen Free™ | 9.7% | (260/2686) | 38.5% | (10/26) |
|  | Excluded Flora and Restricted Flora™ | 1.9% | (29/1537) | 5.9% | (1/17) |
|  | Defined Flora | 0.0% | (0/110) | 0.0% | (0/4) |
|  | Germ free | 0.0% | (0/71) | 0.0% | (0/2) |
| Janvier, Europe | SPF | ND<sup>3</sup> |  | ND<sup>3</sup> |  |
|  | SOPF | 0.0% | (0/303) | 0.0% | (0/6) |
| Envigo (formerly Harlan), Europe | Barrier | ND<sup>3</sup> |  | ND<sup>3</sup> |  |
|  | Isolator, immunocompetent mice | 0.0% | (0/610) | 0.0% | (0/13) |
|  | Isolator, immunodeficient mice | 0.0% | (0/2128) | 0.0% | (0/56) |
| Envigo (formerly Harlan), US | Barrier | 6.0% | (46/761) | 87.5% | (7/8) |
|  | Isolator, immunocompetent mice | 0.0% | (0/85) | 0.0% | (0/3) |
|  | Isolator, immunodeficient mice | 0.0% | (0/2189) | 0.0% | (0/17) |

<sup>1</sup>Standard barrier production colonies (SPF) are designated by the various providers as follows: specific-pathogen-free (Charles River Europe, Janvier), VAF® (virus/antibody free; Charles River, North America), Standard (Jackson, so-called Production, Repository and Breeding Services facilities), Low barrier (Jackson, so-called Research animal facility (RAF)), Murine Pathogen Free (Taconic), and barrier (Envigo). Specific and opportunistic pathogen free (SOPF) colonies are

As a standard procedure at all vendors, laboratory mice colonies in new barrier rooms are set up with *S. aureus*-free breeders. Therefore, *S. aureus* must be accidentally introduced into some colonies at a later stage. Retrospective data obtained from five SPF barriers of Charles River showed that it can take between 10 months and four years until an *S. aureus*-negative SPF barrier turns *S. aureus*-positive (Table S1).

### *S. aureus* efficiently transmits from parents to offspring

To examine *S. aureus* transmission, we selected one *S. aureus* negative and three *S. aureus* positive C57BL/6 breeding pairs and determined gastrointestinal *S. aureus* colonization of the offspring at weaning as well as 4, 8, 12, and 16 weeks later. Notably, *S. aureus* was rapidly and very efficiently transmitted from parents to offspring. All young mice became colonized, most by weaning, and *S. aureus* then persisted in the gut of most mice throughout the period of study (Fig 1). Hence, early exposure to *S. aureus* leads to persistent colonization in laboratory mice. In contrast, the offspring of *S. aureus*-negative mice remained negative. This suggests that once introduced into a facility, *S. aureus* will be readily transmitted to the offspring of colonized breeding pairs but not necessarily between cages.

**Figure 1:**
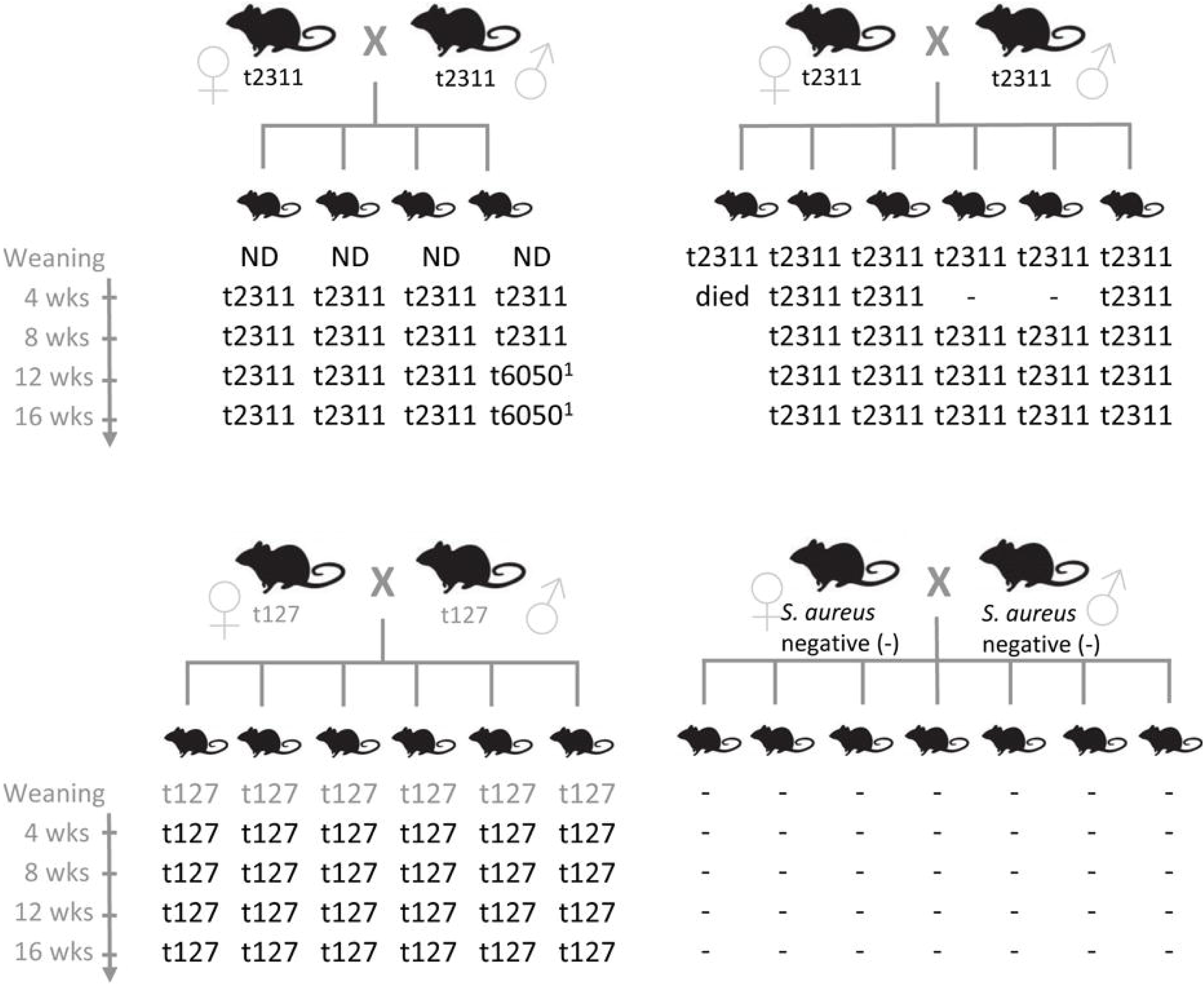
*S. aureus* is efficiently transmitted from murine parents to their offspring. *S. aureus* isolates were obtained from individual stool samples of breeders and their offspring at weaning (20-23 days after birth) as well as 4, 8, 12 and 16 weeks later. Clonal relationship of the *S. aureus* isolates was determined by *spa* typing. Grey *spa* types were derived from a pooled stool sample from either parental mice or offspring. *Spa* types t2133 and t6050 are closely related (both CC88). *Spa* type tl27 belongs to CC1. Key: –, *S. aureus* negative; ND, not determined.

### CC88 is the dominant *S. aureus* lineage in laboratory mice

To determine whether laboratory mice are colonized with typical human isolates or mouse-adapted strains such as JSNZ [10], we characterized a total of 99 *S. aureus* isolates from laboratory mice. *Spa* genotyping was employed to resolve the population structure and compare it to a well-characterized collection of 107 nasal isolates from healthy *S. aureus* carriers from Northern Germany [23]. More than half (54/99) of the murine strains belong to CC88 and were therefore closely related to JSNZ (Fig 2, Table 2). In contrast, we did not detect CC88 among human colonizing strains. Common human lineages (CC5, 8, 12, 15, 25, and 30) accounted for only 37.4% (37/99) of the murine isolates (Fig 2). The livestock-associated lineages CC72 (n=1) and CC188 (n=6) were also detected among the murine isolates [20, 28]. CC88 strains were already present in samples obtained in 2004, suggesting that this lineage is a long-standing companion of laboratory mice (Table 2). Moreover, CC88 strains were widely distributed as these strains were submitted by various customers (pharmaceutical industry, academia, and vendors) from the US, Canada and Japan. Apart from CC88, CC15 isolates were frequent in the sampling cohort, which also originated from various customers and several countries (US, Canada, Great Britain) (Table 2). The described *S. aureus* strain spectrum was also represented in the animal facility of University Medicine Greifswald (Table S3). None of the local animal caretakers carried an animal clone confirming that transmission between humans and mice is rare.

**Figure 2:**
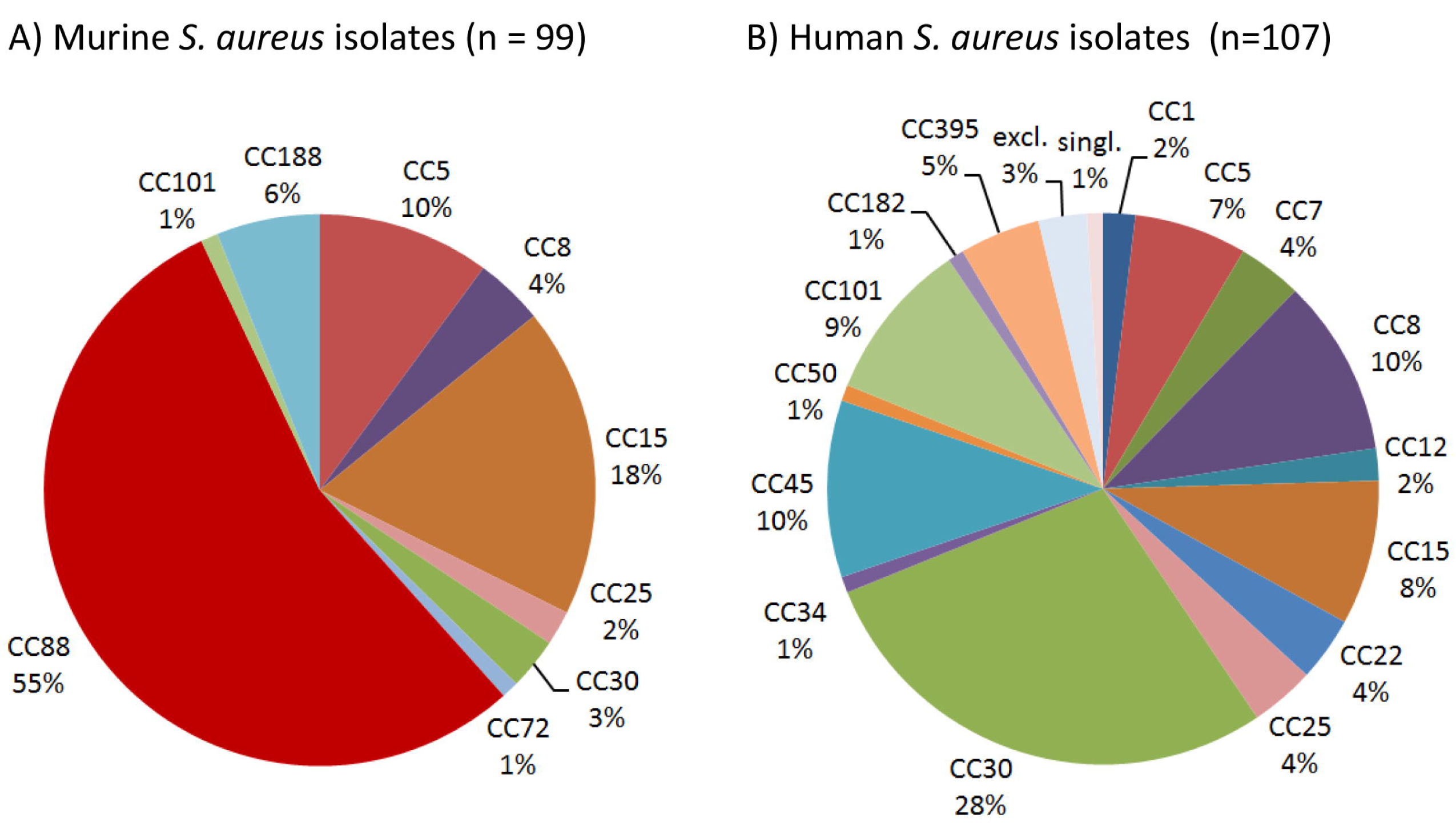
CC88 is common among murine but not human *S. aureus* strains. *Spa* genotyping was employed to resolve the population structure of murine *S. aureus* isolates and compare it to a collection of 107 human *S. aureus* isolates. *Spa* types were clustered into MLST-derived clonal complexes. MLST-CC clustering of the human nasal *S. aureus* isolates (n=107), which has been previously published [23], was refined based on the current SpaServer database. While CC88 was the dominant murine lineage, it was absent from the studied human collection.

**Table 2:**
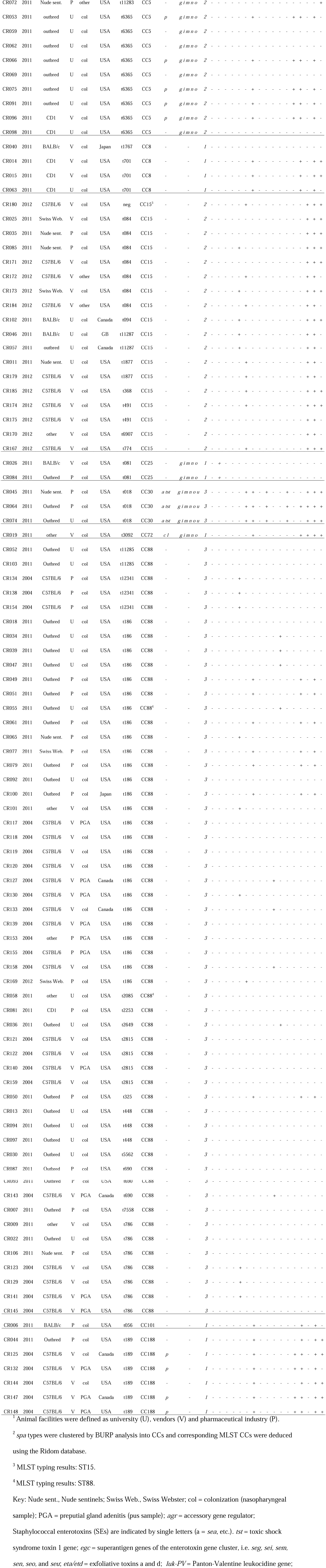
Genotype, virulence genes, phage patterns and ampicillin resistance of *S. aureus* isolates from laboratory mice.

### S. *aureus* isolates from laboratory mice are adapted to their murine host

To investigate host adaptation, we screened the murine *S. aureus* isolates for bacteriophages, MGE-encoded superantigens (SAg), ampicillin resistance and pro-coagulatory activity. A collection of 107 nasal isolates from healthy *S. aureus* carriers (Table S2), supplemented by a total of 24 human CC88 strains from around the globe (Table S4), served as control [23]. Firstly, we screened all murine strains for bacteriophages and the Sa3int phage-encoded human-specific IEC genes. The most prevalent phage families among human isolates, Sa2int and Sa3int phages, were rarely found in murine isolates (Sa2int: 33.6% vs 12.1%, p<0.001; Sa3int: 79.4% vs 26.3%, p<0.001) (Tables 2 and S2). Unexpectedly, all murine and human CC15 isolates harbored *chp* and *scn* but lacked the *Sa3int* gene. These strains carry immobilized remnants of the *Sa3int* phage including the IEC genes *scn* and *chp* in the bacterial genome (S1 Fig). As MGEs are linked to *S. aureus* clonal lineages, we also stratified the prevalence Sa3int phages by CC (Table 3). Again, only 12.9% of murine CC88 strains carried IEC-encoding Sa3int phages compared to 100% of the human CC88 isolates.

**Table 3:** Prevalence of IEC-encoding Sa3int phages, MGE-encoded superantigen genes and ampicillin resistance in murine and human isolates.

|  |  | % positive murine<br>*S. aureus* isolates<br>(absolute No.) | % positive human<br>*S. aureus* isolates<br>(absolute No.) | p value |
| --- | --- | --- | --- | --- |
| IEC-encoding Sa3int<br>phages | CC5 | 50.0% (5/10) | 85.7% (6/7) | n.s. |
|  | CC8 | 75.0% (3/4) | 81.8% (9/11) | n.s. |
|  | CC15<sup>1</sup> | 0.0% (0/18) | 0.0% (0/9) | n.s. |
|  | CC88 | 12.9% (7/54) | 100.0% (24/24) | p<0.001 |
|  | all strains | 26.3% (26/99) | 79.4% (104/131) | p<0.001 |
| MGE-encoded SAg genes | CC5 | 50.0% (5/10) | 57.1% (4/7) | n.s. |
|  | CC8 | 0.0% (0/4) | 100.0% (11/11) | p<0.001 |
|  | CC15 | 0.0% (0/18) | 0.0% (0/9) | n.s. |
|  | CC88 | 0.0% (0/54) | 45.8% (11/24) | p<0.001 |
|  | all strains | 13.1% (13/99) | 59.5% (78/131) | p<0.001 |
| Ampicillin resistance | CC5 | 10.0% (1/10) | 85.7% (6/7) | p<0.01 |
|  | CC8 | 50.0% (2/4) | 81.8% (9/11) | p<0.05 |
|  | CC15 | 55.6% (10/18) | 77.8% (7/9) | n.s. |
|  | CC88 | 0.0% (0/54) | 54.0% (16/24) | p<0.001 |
|  | all strains | 22.2% (22/99) | 66.4% (87/131) | p<0.001 |

<sup>1</sup> All CC15 isolates carry immobilized remnants of the Sa3int phage with most of the phage genome including the integrase missing. The IEC locus, however, is intact (see also Figure S1).
Key: IEC, immune evasion cluster encoding staphylokinase, staphylococcal complement inhibitor, chemotaxis inhibitory protein of *S. aureus*, as well as staphylococcal enterotoxin A or P; MGE, mobile genetic element.

Secondly, we investigated MGE-encoded SAg, because these toxins act on murine T cells with much lower potency than on the human counterparts [29]. MGE-encoded SAgs were detected in 59.5% of the human isolates but only in 13.1% of murine *S. aureus* strains (Table 3) [23]. Mice were typically colonized either with SAg-negative lineages (CC15, CC101) or with SAg-negative variants of lineages harbouring MGE-encoded SAg genes in human isolates (CC88, CC8). For example, 11/24 human CC88 strains were SAg-positive, whereas all 54 murine CC88 isolates were SAg-negative (Tables 2, 3, and S2).

Thirdly, we screened isolates for ampicillin resistance, a common feature of human *S. aureus* strains [13]. Of note, only 22.2% of murine strains were resistant to ampicillin compared to 66.4% of the human isolates (Table 3). All murine isolates were *mecA*-negative.

Finally, we investigated whether murine CC88 *S. aureus* strains are superior to human CC88 strains in coagulating murine plasma. In general, murine plasma coagulated more slowly than human plasma. During the first 4 hours, however, coagulation of murine plasma was more advanced after incubation with murine than with human CC88 isolates (Fig 3). This suggests that murine CC88 *S. aureus* isolates have evolved means to specifically modulate the murine coagulation system.

**Figure 3:**
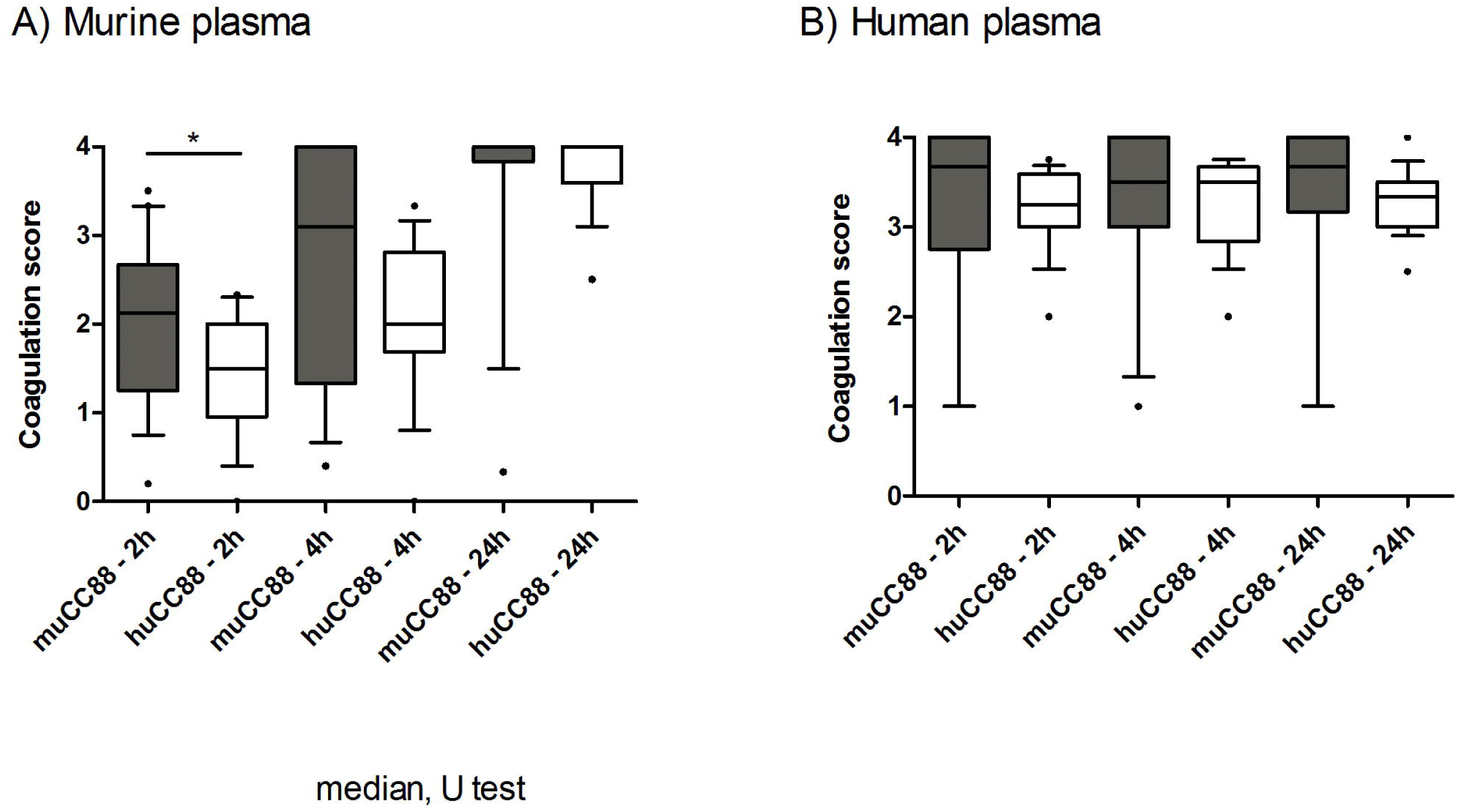
Murine CC88 isolates coagulate murine plasma faster than human CC88 isolates. Murine (n=19) and human (n=17) CC88 isolates were compared in their ability to coagulate murine (A) or human (B) plasma. 500 μl plasma were inoculated with 2.5×10^9^ CFU *S. aureus* and coagulation was visually assessed after 2h, 4h and 24h. At least three technical replicates were performed for each sample, and the average values are depicted as box plots. Boxes indicate the median as well as the 25-75 percentiles; whiskers illustrate the 10-90 percentiles. Statistics: Mann Whitney U test. Key: muCC88, murine CC88 strains; huCC88, human CC88 strains.

### Colonized laboratory mice mount a systemic immune response to *S. aureus*

We have previously reported that symptom-free human *S. aureus* carriers raise a strong serum IgG response against their colonizing strain [30]. To test whether this happens in laboratory mice, we compared antibody profiles of *S. aureus*-free mice with those of symptom-free colonized mice and mice with spontaneous PGA. Animals were derived from two Charles River breeding facilities, Kingston and Hollister, and naturally colonized or infected with strains of the CC88 or CC1 lineage, respectively (Table S5). Importantly, colonized mice showed a significant systemic IgG response against extracellular staphylococcal proteins, whereas SOPF mice were immunologically naïve (Fig S2).

To characterize the induced antibody response in more detail, we quantified serum IgG binding to a panel of 58 recombinant *S. aureus* proteins. A 2-fold rise in antibody titers in exposed vs. SOPF mice was considered as biologically relevant. A total of seven *S. aureus* proteins were recognized by all *S. aureus-*exposed mice: Plc, Atl, HlgB, HlgC, Hlb, SplD, and PknB (Fig 4, Table S6). Moreover, colonized mice were primed against vaccine candidates from previous or ongoing human clinical trials (ClfB, Cna, Hla, IsdB, SdrE, and SdrG) [2]. Notably, colonized mice from both facilities showed more than 100-fold higher Plc-specific antibody titers than SOPF mice, which makes Plc a suitable candidate for a serological screening assay for *S. aureus* exposure (S2 Fig).

**Figure 4:**
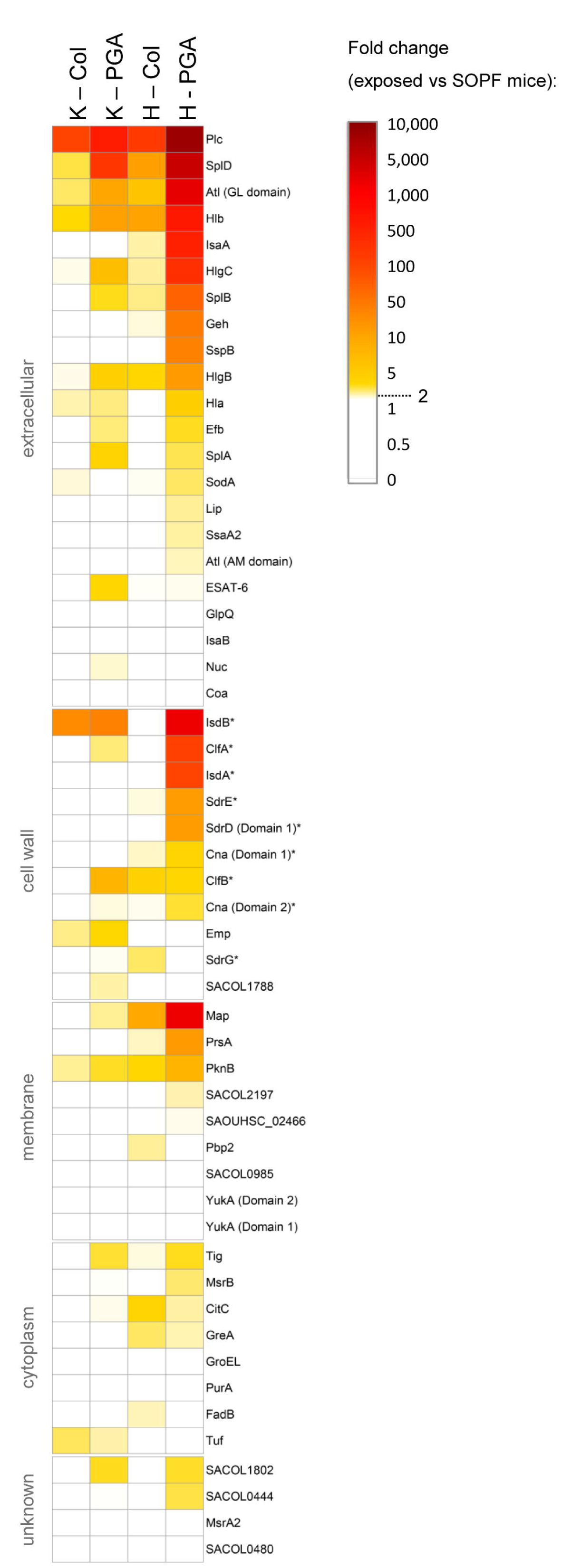
*S. aureus*-exposed laboratory mice mount a serum IgG response against a range of *S. aureus* proteins, including vaccine candidates. A) Antigen-specific murine serum IgG was measured using FlexMap^®^ technology. We measured the strain-specific antibody response against 58 recombinant *S. aureus* antigens in colonized symptom-free mice as well as in mice with spontaneous infections (PGA) from Charles River breeding facilities at Kingston and Hollister. SOPF mice served as negative controls. Group size was n = 5. Mice from Kingston were exposed to a CC88 isolate, while mice from Hollister were exposed to a CC1 isolate. Fold-changes of exposed mice as compared to the SOPF controls are depicted. Values below 2.0 are displayed in white. B) Dot plot of the phospholipase C-specific serum IgG response as determined FlexMap^®^ technology. MFI values for three SOPF sera were below the limit of detection (<1.47) and are thus not displayed on the graph. Key: Col, colonization; PGA, preputial gland adenitis; K, Kingston (mice were exposed to CC88 isolates); H, Hollister (mice were exposed to CC1 isolates); MFI = Mean fluorescence intensity.

Antibody titers against numerous antigens were higher in infected mice from Hollister (CC1) than in their counterparts from Kingston (CC88) (Fig 4, Table S6). Since all tested antigens are encoded in the CC88 genome (unpublished data), the more pronounced immune response might be due to differences in the in vivo behavior of CC1 and CC88 isolates.

## Discussion

Many *S. aureus* researchers doubt that mice are a suitable infection model for *S. aureus* research because mice are not considered to be natural hosts of these bacteria [3-6]. In this study, we report that SPF laboratory mice from all the main vendors are colonized with *S. aureus* at varying rates. This is not surprising because *S. aureus* is considered an opportunistic pathogen in mice and not routinely excluded from SPF barrier rooms. Of note, vendors respond very differently to the presence of opportunistic agents. Some vendors tolerate these organisms in SPF colonies, whereas other vendors test and cull mice to stop their spread. In Europe, FELASA recommendations do not require vendors to publish *S. aureus* test results from SPF mice [21]. Instead, European researchers must approach each company for information about the *S. aureus* colonization status of their mouse strain of interest. Alternatively, SOPF animals from all vendors are by definition *S. aureus-*free.

We demonstrated that *S. aureus* is readily transmitted from parents to offspring, who become persistently colonized. Retrospective data on *S. aureus* prevalence in SPF barriers at Charles River showed that mouse colonies are not able to clear *S. aureus* once it has been introduced into a colony. These findings imply that *S. aureus* is maintained by efficient vertical transmission, and that early exposure may be decisive for long-term colonization possibly due to early occupation of niches in the gut and nose. In contrast, human to mouse transmission or acquisition from the environment appears to be much less frequent. In this study, the *S. aureus* strains predominating in animal facilities (CC88 lineage) are rarely found in the human population. Moreover, we observed no concordance of murine and human *S. aureus* isolates at a local university-associated animal facility. This contrasts studies from the 1970s and 80s, which reported similar phage patterns between isolates from caretakers and infected mice [11, 12]. This suggests that nowadays a host jump is efficiently limited by using protected cage changing stations and isolation housing, performing regular microbiological screenings and adhering to strict hygiene practices [8].

The predominant lineage among the murine isolates was CC88, which is known for causing an outbreak of PGA at an animal facility at Auckland University [10]. Remarkably, CC88 strains (1) have been persisting in the Charles River facilities for at least a decade, (2) were efficiently transmitted from murine parents to offspring, and (3) were detected in animal facilities around the globe (e.g. New Zealand, US and Germany). This strongly suggests that laboratory mice are a major reservoir for this CC88 sub-lineage. To our best knowledge, CC88 strains have never been detected in animals. However, outbreaks of CC88 CA-MRSA infections in humans have been reported from Asia and Africa [14, 15]. Hence, the murine CC88 population might be the result of a human-to-mouse host jump, followed by genetic adaptation to the murine host.

Our comparisons of human and murine *S. aureus* strains suggest that adaptation to the murine host could involve the loss of Sa3int phages, SAg-encoding MGEs and antibiotic resistance genes. Sa3int phages encode immune evasion factors but also destroy the *hlb* gene upon integration into the bacterial chromosome (Fig S1). In the murine system, the IEC-encoded factors show no or negligible activity [29, 31-33], whereas the sphingomyelinase Hlb mediates hemolysis of ruminant erythrocytes and is an important virulence factor in mouse infection models [34]. Thus, in the murine host, the disadvantage due to loss of Hlb clearly outweighs the benefits conferred by the IEC-encoded factors. The absence of Sa3int phages, which is accompanied by ß-hemolysis, is indeed a common feature of animal-adapted strains [20, 35].

Penicillin resistance, mediated by the production of the *blaZ-*encoded ß-lactamase, was rare among murine isolates. Maybe the host jump occurred before the spread of ampicillin resistance in the human *S. aureus* population in the 1950s [36]. Alternatively, the ß-lactamase-encoding plasmids or transposons might have been lost following the host jump [37]. Commercial vendors do not use antibiotics in barriers, thereby removing the selective pressure for maintaining the resistance gene.

The ability to coagulate plasma is an essential virulence trait of *S. aureus,* and some pro- or anti-coagulatory factors are host-specific [38]. This makes it a strong and easily addressable indicator of host-adaptation. Here, we show that murine CC88 isolates coagulate murine plasma faster than matched human CC88 isolates. In line with our data, Viana *et al.* reported that ruminant strains, but not human isolates, had the capacity to stimulate clotting of ruminant plasma [38]. In some ruminant strains, this effect is based on an MGE-encoded paralogue of the van Willebrand factor binding protein. Whether the enhanced coagulation of murine plasma is mediated by a similar mechanism or rather by the acquisition of MGEs with novel pro-coagulatory factors will be addressed by whole genome sequencing of the isolates. The frequent colonization of laboratory mice with *S. aureus* is of concern because it could significantly affect *S. aureus* infection and vaccination studies. Here, we show that colonized SPF mice mount a systemic antibody response against a panel of *S. aureus* proteins, including numerous vaccine candidates from past or current clinical trials [2]. In contrast, commensal gut bacteria usually do not elicit a systemic IgG response but a local IgA response. This suggests that *S. aureus* is more aggressive than gut commensals and probably induces minor subclinical *S. aureus* infection in mice which then trigger a systemic immune response as has been suggested for humans [39].

Importantly, immune priming of laboratory mice before experimental infection or vaccination may strongly influence the outcome [27, 39]. Our data suggests that unrecognized *S. aureus* colonization of mice may be a significant confounder in experimental infection and vaccination studies, especially if prior exposure to *S. aureus* is variable. We strongly encourage researchers to ensure that they work with either *S. aureus-*free or consistently primed mice.

Since *S*. *aureus* screening results from commercial vendors are not available for SPF animals from European facilities, a microbiological or serological screening assay will be of advantage. Nose homogenates or stool samples are suitable for microbiological screenings, but the available selective media are error prone. In contrast, Plc, for which we observed a more than 100-fold increase in antibody titers in *S. aureus-*exposed mice, could be a robust and sensitive marker for a serological screening assay for *S. aureus* exposure.

Robust and clinically relevant infection models are mandatory for the development of new strategies to prevent or treat *S. aureus* infection. Our data shows that laboratory mice are better models for colonization and infection studies than previously assumed. Just like humans, laboratory mice are persistently colonized with *S. aureus,* mount a systemic immune response to colonization, and frequently suffer from abscess formation. Moreover, the use of mouse-adapted *S. aureus* strains in their natural host – the mouse - promises to provide a more physiological model for studying *S. aureus* host interaction and testing novel therapeutics. Such strains can show better fitness and virulence in mouse colonization and infection models, respectively [10]. However, one has to keep in mind that mice and mouse-adapted strains are not suited to study the effect of human-specific virulence factors, such as PVL and SAgs [40].

## Confict of interests

S. Monecke is an employee of Alere Technologies, a company that manufactures microarrays. The company had no influence on the study design, experiments, data interpretation, and publication.

K. Pritchett-Corning and R. Fister were previously employed at Charles River, USA. The company had no influence on the study design, experiments, data interpretation, and publication.

## Presentation of data

A part of the data has been previously presented as a poster at the International Symposium on Staphylococci and Staphylococcal Diseases in Chicago, USA in August 2014 and at the 3^rd^ International Conference on the Pathophysiology of Staphylococci in Tübingen, Germany in September 2016.

## Acknowledgement

We would like to thank Anne Morin, Erika Friebe, Stefanie Förster, Katrin Schmoeckel, and Fawaz Al Sholui for technical support, Stefan Weiß for support in data visualization, and Johannes Dick and Dina Raafat for helpful comments on the manuscript.

## Supporting information

### Text S1. Full-length Materials and Methods

**S1 Fig. CC15 strains carry remnants of the *hlb*-integrating Sa3int phage.** A) Schematic illustration of the site-specific integration of phages of the Sa3int family. Side-specific integration is mediated by attp sites (orange) which are homologous to sequences within the *hlb* gene (brown). Integration of Sa3int phages leads to insertion of the phage genome including the immune evasion gene cluster (IEC; green) into the bacterial chromosome and to a disruption of the *hlb* gene. In contrast, *S. aureus* strains lacking Sa3int phages have an intact *hlb* gene. B) Comparison of whole genome sequences of Newman (CC8), VCU006 and CIG93 (both CC15). CC15 strains carry immobilized remnants of the phage including the IEC genes *scn* and *chp* (green). Since 35.8kb of the phage genome, including the 3' attp side (orange) and integrase (red) are deleted, these phage remnants cannot be mobilized any more.

**S2 Fig. *S. aureus-*exposed laboratory mice mount a serum IgG response against extracellular *S. aureus* proteins.** We measured the strain-specific antibody response in colonized symptom-free mice as well as in mice with spontaneous infections (PGA) from Charles River breeding facilities at Kingston and Hollister. SOPF mice served as negative controls. Group size was n = 5. Mice from Kingston were exposed to a CC88 isolate closely related to JSNZ, while mice from Hollister were exposed to a CC1 isolate. A, B) Serum IgG binding to extracellular proteins of *S. aureus* JSNZD*spa* (A) or the CC1 isolate PGA12D*spa* (B) was determined by ELISA. One out of two very similar experiments is depicted. Graphs show the median for each group. Groups were compared using the Mann-Whitney test with a Bonferroni correction. C) Dot plot of the phospholipase C-specific serum IgG response as determined FlexMap^®^ technology. MFI values for three SOPF sera were below the limit of detection (<1.47) and are thus not displayed on the graph. Key: SOPF, specific opportunistic pathogen free; Col, colonization; PGA, preputial gland adenitis.

**S1 Table. Percentage of *S*. *aureus*-positive mice and barriers in different breeding facilities, sorted by vendor and barrier status (cumulative data extracted from publicly available health reports in November 2016).**

^1^Standard barrier production colonies (SPF) are designated by the various providers as follows: specific-pathogen-free (Charles River Europe, Janvier), VAF^®^ (virus/antibody free; Charles River, North America), Standard (Jackson, so-called Production, Repository and Breeding Services facilities), Low barrier (Jackson, so-called Research animal facility (RAF)), Murine Pathogen Free (Taconic), and barrier (Envigo). Specific and opportunistic pathogen free (SOPF) colonies are designated as follows: SOPF (Charles River Europe, Janvier), Elite^®^ (Charles River, North America), intermediate barrier and high barrier (Jackson), Excluded Flora/Restricted Flora (Taconic), and Isolator (Envigo). Each vendor has specific definitions of its barriers. For details refer to the vendor's webside.

^2^Cumulative data comprising the last 18 months (Charles River, Janvier, and Envigo), 12 months (The Jackson laboratories) or 6 months (Taconic). Envigo reported the number of positive isolators (not barriers).

^3^*S. aureus* not reported based on FELASA recommendations (2002 and 2014)

^4^According to the company, a test and cull effort was initiated to eliminate *S. aureus* from this barrier. This investigation has been completed, and no additional positives have been found.

**S2 Table. Genotype, virulence genes, phage patterns and ampicillin resistance of *S. aureus* isolates from laboratory mice.**

^1^ Animal facilities were defined as university (U), vendors (V) and pharmaceutical industry (P).

^2^*spa* types were clustered by BURP analysis into CCs and corresponding MLST CCs were deduced using the Ridom database.

^3^ MLST typing results: ST15.

^4^ MLST typing results: ST88.

Key: Nude sent., Nude sentinels; Swiss Web., Swiss Webster; col = colonization (nasopharyngeal sample); PGA = preputial gland adenitis (pus sample); *agr* = accessory gene regulator; Staphylococcal enterotoxins (SEs) are indicated by single letters (a = *sea,* etc.). *tst* = toxic shock syndrome toxin 1 gene; *egc* = superantigen genes of the enterotoxin gene cluster, i.e. *seg, sei, sem, sen, seo,* and *seu; eta/etd* = exfoliative toxins a and d; *luk-PV* = Panton-Valentine leukocidine gene; *Sa1int* = *S. aureus* integrase type 1; *sak =* Staphylokinase gene, *chp* = gene encoding the chemotaxis inhibitory protein; *scn* = staphylococcal complement inhibitor gene; AmpR = ampicillin resistance

S3 Table. Genotype, virulence genes and phage patterns of colonizing *S. aureus* isolates from randomly sampled mice and their animal care takers at a university breeding facility.

S4 Table. Genotype, virulence genes, phage patterns and ampicillin resistance of colonizing *S. aureus* isolates from healthy blood donors.

S5 Table. Genotype, virulence genes, phage patterns and ampicillin resistance of human CC88 *S. aureus* isolates.

S6 Table. Binding of murine IgG serum antibodies to recombinant *S. aureus* proteins using the FlexMAP system.

## References

1. World Health Organization. Antimicrobial resistance: global report on surveillance. In: Press W, ed. Geneva, Switzerland: World Health Organization, 2014.

2. Fowler VG, Jr., Proctor RA. Where does a Staphylococcus aureus vaccine stand? Clin Microbiol Infect 2014; 20 Suppl 5:66-75.

3. Mulcahy ME, Geoghegan JA, Monk IR, et al. Nasal colonisation by Staphylococcus aureus depends upon clumping factor B binding to the squamous epithelial cell envelope protein loricrin. PLoS Pathog 2012; 8:e1003092.

4. Capparelli R, Nocerino N, Medaglia C, Blaiotta G, Bonelli P, Iannelli D. The Staphylococcus aureus peptidoglycan protects mice against the pathogen and eradicates experimentally induced infection. PLoS One 2011; 6:e28377.

5. McCarthy AJ, Lindsay JA. Genetic variation in Staphylococcus aureus surface and immune evasion genes is lineage associated: implications for vaccine design and host-pathogen interactions. BMC Microbiol 2010; 10:173.

6. Cuny C, Friedrich A, Kozytska S, et al. Emergence of methicillin-resistant Staphylococcus aureus (MRSA) in different animal species. Int J Med Microbiol 2010; 300:109-17.

7. Kiser KB, Cantey-Kiser JM, Lee JC. Development and Characterization of a *Staphylococcus aureus* Nasal Colonization Model in Mice. Infect Immun 1999; 67:5001-6.

8. Baker DG. Natural pathogens of laboratory mice. Their effects on research. Vol. 1. Washington: American Society for Microbiology (ASM) Press, 2003.

9. Pritchett-Corning KR, Cosentino J, Clifford CB. Contemporary prevalence of infectious agents in laboratory mice and rats. Lab Anim 2009; 43:165-73.

10. Holtfreter S, Radcliff FJ, Grumann D, et al. Characterization of a mouse-adapted Staphylococcus aureus strain. PLoS One 2013; 8:e71142.

11. Blackmore DK, Francis RA. The apparent transmission of staphylococci of human origin to laboratory animals. J Comp Pathol 1970; 80:645-51.

12. Needham JR, Cooper JE. Bulbourethral gland infections in mice associated with Staphylococcus aureus. Lab Anim 1976; 10:311-5.

13. Holtfreter S, Grumann D, Balau V, et al. Molecular epidemiology of Staphylococcus aureus in the general population in Northeast Germany - results of the Study of Health in Pomerania (SHIP-TREND-0). J Clin Microbiol 2016.

14. Ghebremedhin B, Olugbosi MO, Raji AM, et al. Emergence of a community-associated methicillin-resistant Staphylococcus aureus strain with a unique resistance profile in Southwest Nigeria. J Clin Microbiol 2009; 47:2975-80.

15. Zhang W, Shen X, Zhang H, et al. Molecular epidemiological analysis of methicillin-resistant Staphylococcus aureus isolates from Chinese pediatric patients. Eur J Clin Microbiol Infect Dis 2009; 28:861-4.

16. Gorwitz RJ, Kruszon-Moran D, McAllister SK, et al. Changes in the prevalence of nasal colonization with Staphylococcus aureus in the United States, 2001-2004. J Infect Dis 2008; 197:1226-34.

17. Guinane CM, Ben Zakour NL, Tormo-Mas MA, et al. Evolutionary genomics of Staphylococcus aureus reveals insights into the origin and molecular basis of ruminant host adaptation. Genome Biol Evol 2010; 2:454-66.

18. Herron-Olson L, Fitzgerald JR, Musser JM, Kapur V. Molecular correlates of host specialization in Staphylococcus aureus. PLoS One 2007; 2:e1120.

19. van Wamel WJ, Rooijakkers SH, Ruyken M, van Kessel KP, van Strijp JA. The innate immune modulators staphylococcal complement inhibitor and chemotaxis inhibitory protein of *Staphylococcus aureus* are located on beta-hemolysin-converting bacteriophages. J Bacteriol 2006; 188:1310-5.

20. Sung JM, Lloyd DH, Lindsay JA. Staphylococcus aureus host specificity: comparative genomics of human versus animal isolates by multi-strain microarray. Microbiology 2008; 154:1949-59.

21. Mahler Convenor M, Berard M, Feinstein R, et al. FELASA recommendations for the health monitoring of mouse, rat, hamster, guinea pig and rabbit colonies in breeding and experimental units. Lab Anim 2014.

22. Goerke C, Pantucek R, Holtfreter S, et al. Diversity of prophages in dominant Staphylococcus aureus clonal lineages. J Bacteriol 2009; 191:3462-8.

23. Holtfreter S, Grumann D, Schmudde M, et al. Clonal distribution of superantigen genes in clinical Staphylococcus aureus isolates. J Clin Microbiol 2007; 45:2669-80.

24. Enright MC, Day NP, Davies CE, Peacock SJ, Spratt BG. Multilocus sequence typing for characterization of methicillin-resistant and methicillin-susceptible clones of *Staphylococcus aureus.* J Clin Microbiol 2000; 38:1008-15.

25. Harmsen D, Claus H, Witte W, Rothganger J, Turnwald D, Vogel U. Typing of methicillin-resistant Staphylococcus aureus in a university hospital setting by using novel software for spa repeat determination and database management. J Clin Microbiol 2003; 41:5442-8.

26. Sperber WH, Tatini SR. Interpretation of the tube coagulase test for identification of Staphylococcus aureus. Applied microbiology 1975; 29:502-5.

27. Stentzel S, Sundaramoorthy N, Michalik S, et al. Specific serum IgG at diagnosis of Staphylococcus aureus bloodstream invasion is correlated with disease progression. Journal of proteomics 2015; 128:1-7.

28. McCarthy AJ, Lindsay JA, Loeffler A. Are all meticillin-resistant Staphylococcus aureus (MRSA) equal in all hosts? Epidemiological and genetic comparison between animal and human MRSA. Veterinary dermatology 2012; 23:267-75, e53-4.

29. Holtfreter S, Broker BM. Staphylococcal superantigens: do they play a role in sepsis? Arch Immunol Ther Exp (Warsz) 2005; 53:13-27.

30. Holtfreter S, Roschack K, Eichler P, et al. Staphylococcus aureus carriers neutralize superantigens by antibodies specific for their colonizing strain: a potential explanation for their improved prognosis in severe sepsis. J Infect Dis 2006; 193:1275-8.

31. Rooijakkers SH, Ruyken M, Roos A, et al. Immune evasion by a staphylococcal complement inhibitor that acts on C3 convertases. Nat Immunol 2005; 6:920-7.

32. de Haas CJC, Veldkamp KE, Peschel A, et al. Chemotaxis Inhibitory Protein of *Staphylococcus aureus,* a Bacterial Antiinflammatory Agent. J Exp Med 2004; 199:687-95.

33. Gladysheva IP, Turner RB, Sazonova IY, Liu L, Reed GL. Coevolutionary patterns in plasminogen activation. Proc Natl Acad Sci U S A 2003; 100:9168-72.

34. Katayama Y, Baba T, Sekine M, Fukuda M, Hiramatsu K. Beta-hemolysin promotes skin colonization by Staphylococcus aureus. J Bacteriol 2013; 195:1194-203.

35. Markham NP, Markham JG. Staphylococci in man and animals. Distribution and characteristics of strains. J Comp Pathol 1966; 76:49-56.

36. Chambers HF, Deleo FR. Waves of resistance: Staphylococcus aureus in the antibiotic era. Nat Rev Microbiol 2009; 7:629-41.

37. Lyon BR, Skurray R. Antimicrobial resistance of Staphylococcus aureus: genetic basis. Microbiol Rev 1987; 51:88-134.

38. Viana D, Blanco J, Tormo-Mas MA, et al. Adaptation of Staphylococcus aureus to ruminant and equine hosts involves SaPI-carried variants of von Willebrand factor-binding protein. Mol Microbiol 2010; 77:1583-94.

39. Broker BM, Holtfreter S, Bekeredjian-Ding I. Immune control of Staphylococcus aureus - regulation and counter-regulation of the adaptive immune response. Int J Med Microbiol 2014; 304:204-14.

40. Loffler B, Hussain M, Grundmeier M, et al. Staphylococcus aureus panton-valentine leukocidin is a very potent cytotoxic factor for human neutrophils. PLoS Pathog 2010; 6:e1000715.

